# Fast structural search for classification of gut bacterial mucin O-glycan degrading enzymes

**DOI:** 10.64898/2026.02.17.706273

**Authors:** Mert Erden, Tyler Schult, Karin Yanagi, Jugal Kishore Sahoo, David L. Kaplan, Lenore J. Cowen, Kyongbum Lee

## Abstract

The Enzyme Commission (EC) numbering scheme provides a hierarchical way to classify enzymes according to their catalytic functions. While recent protein language model (PLM) based approaches like CLEAN and ProteInter have improved sequence-based EC number prediction, they struggle with fine-grained classification at the deepest hierarchical level. Structure-based approaches for grouping similar proteins using alignment tools excel at finding proteins that share overall global structure, but suffer from high false positive rates when classifying proteins that are globally structurally similar but functional differentiation depends on a localized region. This problem is particularly relevant to EC number prediction, as enzymatic function depends on its catalytic domain, which is a relatively small, specific region of the protein. We introduce Deep Enzyme Function Transfer (DEFT) that harmonizes sequence- and structure-based approaches through the key insight that PLM based annotations of the first two EC number hierarchy levels vastly reduce false positives that are likely to show in purely structure-based EC number prediction. Given an enzyme of interest, DEFT first uses a PLM based method to assign the first two levels of the enzyme’s EC number, and then uses a structure-based method to predict the remaining two levels of the EC number. Using benchmarking datasets, we demonstrate that DEFT achieves superior accuracy compared with current state-of-the-art tools for EC number prediction. Furthermore we show that DEFT’s computational efficiency enables high-throughput, genome-wide annotations of total enzyme repertoires in organisms. We illustrate this capability by experimentally validating DEFT predicted glycoside hydrolase (GH) profiles of intestinal mucus associated bacteria.

**Author summary:** Enzymes are ubiquitous proteins that catalyze chemical reactions of living cells. Enzymes are classified using a hierarchical numbering system called Enzyme Commission (EC) numbers that describe the chemical reactions the enzymes catalyze, from a general reaction type (e.g., breaking bonds, transferring chemical groups, etc.) to more specific aspects such as chemical bonds and substrates involved in the reaction. We present a new machine learning method for predicting EC numbers called Deep Enzyme Function Transfer (DEFT). This method improves on previous methods that use either protein sequence- or three-dimensional (3D) structure-based comparisons between enzymes of known and unknown classification. DEFT combines the strengths of both approaches by first using a protein sequence-based model to predict the general enzyme category and then using protein structure comparisons to predict the finer subcategories. We demonstrate that DEFT achieves superior accuracy compared with current state-of-the-art tools for EC number prediction. We next demonstrate how DEFT’s computational efficiency enables us to perform high-throughput, genome-wide annotations of organisms’ enzyme repertoires. We illustrate this capability by experimentally validating DEFT predicted sugar metabolizing enzyme profiles of intestinal mucus associated bacteria.

## Introduction

Enzymes are the main catalytic units in metabolism. Collectively, the enzymes of an organism describe the biochemical reactions that are possible for the organism since almost all cellular reactions are enzyme-catalyzed. The standard [1] way of describing the catalytic function of an enzyme is to assign an Enzyme Commission (EC) number to the enzyme, which, by construction, is hierarchical. Unlike an accession number in UniProt, which is unique to the corresponding protein, the same EC number can be assigned to two or more proteins if the corresponding enzymes catalyze the same reaction. In addition, a single enzyme can be assigned more than one EC number if it is multifunctional and catalyzes multiple reactions. In some cases, a single EC number represents multiple reactions of a similar type. In other cases, a single EC number can represent two or more distinct reactions because it is associated with an enzyme that is pleiotropic or exhibits substrate promiscuity. The complexity of relationships between enzymes, EC numbers, and reactions [2] has presented challenges in developing efficient algorithms to systematically classify enzymes based on their catalytic functions.

Recently, several deep learning methods have been developed to predict the EC number of an enzyme directly from its amino acid sequence [3–7] but even the best of these methods struggle to correctly assign EC numbers, especially down to the lowest level (fourth digit). When predicting the function of an enzyme, its protein structure adds valuable information because the catalytic mechanism depends on the shape and physicochemical properties of the enzyme’s substrate binding site(s) [8]. Given this, it follows that structure-based methods could help with the EC number prediction task. However, even when an enzyme’s full 3D structure is available from Alphafold2 [9], Alphafold3 [10] or a related protein structure prediction tool (ESMfold [11], OmegaFold [12], etc.), it remains an open question how these structures can be used to effectively predict the enzyme’s EC number without resorting to computationally expensive docking methods. A naive approach is to find an annotated enzyme that is the closest 3D structure match to the enzyme of interest using some global structural alignment method (e.g., TMalign [13]), and then transfer the annotated enzyme’s EC number to the enzyme of interest. However, this approach can produce false positive annotations because enzymes having similar global structures can have distinct catalytic regions due to divergent evolution.

We introduce a new EC number prediction method, Deep Enzyme Function Transfer (DEFT), that combines the advantages of sequence-based deep learning and structure-based search to achieve superior enzyme classification performance. This new method leverages computationally predicted enzyme 3D structure in two ways. First, DEFT uses SaProt [14], a structure-informed protein language model (PLM), to predict the first two EC number levels (class and subclass), for an enzyme of interest. Then, DEFT uses the Foldseek [15] structure-based 3Di string to find the closest annotated structural match having the same first two EC number levels as the enzyme of interest. That matching enzyme’s full EC number (class, subclass, sub-subclass, and serial number) is then transferred to the enzyme of interest to complete the EC number prediction. We benchmark DEFT against several other EC number prediction tools and show that it achieves superior recall and precision scores on the benchmark datasets.

In addition to demonstrating DEFT’s strong performance in annotating individual enzymes, we also show that the method, due to its computational efficiency, enables high-throughput, genome-wide annotations of enzyme repertoires. We evaluated this capability by applying DEFT to the problem of predicting glycoside hydrolase (GH) functions of enzymes in intestinal mucus associated bacteria. The GHs associated with O-glycan degradation are often pleiotropic or exhibit substrate promiscuity [16–18], and span structurally diverse proteins [19, 20]. This has limited computational efforts to identify and annotate these enzymes in different organisms of interest. Applied to a representative set of known mucin grazing and non-grazing gut bacteria, genome-wide scans by DEFT predicted a more extensive set of O-glycan metabolizing GHs for the mucin grazers compared with the non-grazers. We experimentally validated these predictions by culturing the bacteria in media supplemented with various mucins.

Bacteria predicted to have the requisite GHs significantly increased the levels of core and terminal glycan sugars in the culture medium when incubated with mucins, whereas bacteria predicted to lack all or some of the GHs did not show the increases. Together, our results demonstrate that DEFT is a powerful computational method to rapidly and accurately screen whole genomes for metabolic phenotypes of interest.

## Materials and methods

### DEFT

Enzyme annotation by DEFT can be split into three distinct steps: coarse prediction, alignment, and filtering (Fig. 1). We outline each step below.

**Fig 1.**
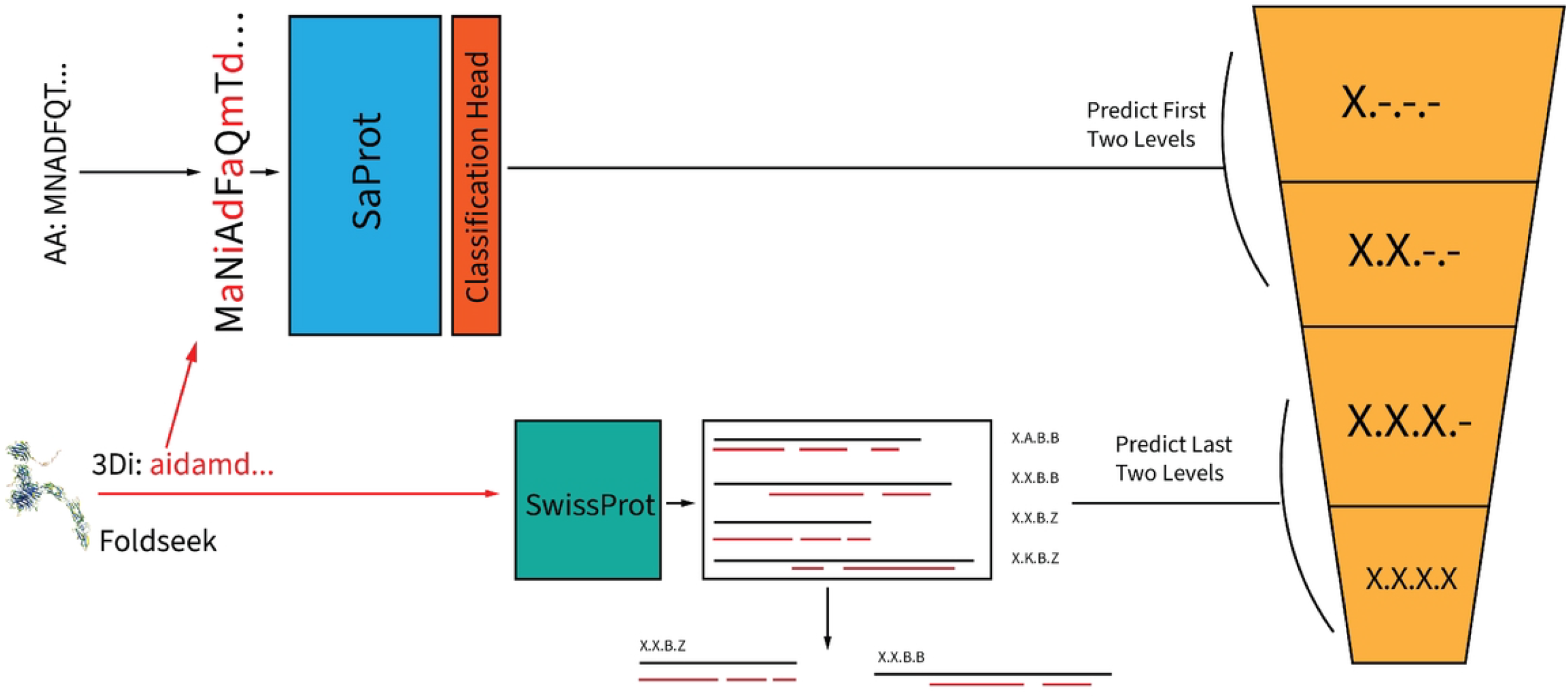
Schematic overview of enzyme classification by DEFT. In step 1, the primary amino acid sequence (AA) and structural representation (3Di) of the enzyme of interest are merged and passed to a fine-tuned protein language model (PLM) to predict the first two levels of the EC number hierarchy. In step 2, the enzyme’s structure is aligned against a reference database. In step 3, the predicted first two digits of the EC number (“prefix”) from the PLM is used to filter out the alignments, where an alignment is kept if the prefix matches the first two digits of the predicted EC number. The final output is a ranked list of EC number predictions for the enzyme of interest sorted by E-values, where the E-value correlates with the probability that the EC number match occurred by chance. See the methods section for details of the E-value calculation.

### Coarse prediction

The EC number system classifies enzymes using a four-level hierarchy based on the reaction they catalyze, with each digit providing increasing specificity. Given the protein sequence and 3D structure of an enzyme of interest, we use the SaProt [14] PLM embedding to learn how to predict the first two EC number digits. The Fine Prediction step, described next, completes the EC number prediction. SaProt’s input alphabet pairs amino acids *A* with structural states *S*, yielding proteins as sequences (*a*_1_*s*_1_, …, *a*_*m*_*s*_*m*_) where *a*_*i*_∈ *A*, ∈*s*_*i*_ *S*. With mask tokens, this creates 441 unique inputs. Fine-tuning uses LoRA [21] and a standard multilayer perceptron classifier trained via cross-entropy loss.

In our coarse classification step, we ignore the hierarchy in the first two EC number levels and instead treat each EC number prefix (through level 2) as a separate class. Thus the coarse classification task assigns each enzyme to one of 78 classes. Freezing the granularity at level 2 and not attempting to predict the entire 4-level EC number as separate stand-alone labels at this stage both reduces the variance that would emerge from training a model with a large number of classes, and allows us to take advantage of the hierarchical EC number structure in the next, fine classification step.

### Fine prediction

Taking the coarse prediction above, DEFT next leverages the hierarchical structure of EC numbers and completes the final two levels using structural alignment. Specifically, DEFT performs a local alignment against a reference database on the 3Di representation of the enzyme returning all sufficiently strong matches, to be filtered in the final step. We utilize Foldseek’s optimized implementation of the Smith-Waterman algorithm [22] for structure matching.

### Filtering and scoring

After the coarse prediction and alignment step, entries with the lowest E-values are selected and their annotations are transferred as the final prediction. In the Results section showing enzyme classification performance on benchmark data, we compare the final prediction to the baseline structural alignment method that predicts all four levels of the EC number solely based on the closest Foldseek match.

To rank DEFT’s predictions we rely on Foldseek E-values, which are analogous to standard sequence alignment E-values but differs in several important ways. Foldseek E-values are defined as follows.

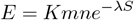

where *S* is a raw alignment score as reported by the alignment algorithm, *λ* and *K* are statistical parameters that control decay rate and densities respectively, and *m* and *n* are the lengths of the aligned sequences. Foldseek [15] dynamically controls the hyperparameters *K* and *λ* and provides a predetermined substitution matrix *S* that is calibrated for the structural alignment task. The resulting Foldseek alignment scores (which we utilize unchanged) are similar to E-values seen in primary sequence alignment, where lower values imply a statistically more likely match, i.e., the probability a random pair of sequences could achieve this alignment is lower. However, because the alignment scores from Foldseek are based on different hyperparameters, they do not have the same distribution as sequence alignment E-values, and a direct numeric comparison cannot be made. Thus, to estimate E-values for each match, Foldseek trains a neural network to predict the mean *µ* and scale parameter *λ* of the extreme value distribution for each query.

### Bacterial strains and cell culture reagents

All chemicals and reagents were purchased from MilliporeSigma (Burlington, MA) or Thermo Fisher Scientific (Waltham, MA) unless otherwise specified. Seven anaerobic bacterial strains were selected for *in vitro* cell culture to experimentally evaluate DEFT’s predictions of mucin O-glycan utilization. The selected strains comprised two known mucin grazers, four non-grazers, and a subspecies that can conditionally degrade mucin O-glycans. The mucin grazers were *Akkermansia muciniphila* ATCC BAA-835 (Am) and *Bacteroides thetaiotaomicron* ATCC 29148 (Bt). The non-grazers were *Lactobacillus plantarum* ATCC 8014 (Lp), *Lactobacillus reuteri* ATCC 23272 (Lr), *Bifidobacterium dentium* ATCC 27678 (Bd), and *Escherichia coli* Nissle (EcN). The conditional mucin-grazer was a strain (ATCC 15707) of *Bifidobacterium longum subsp. longum* (Bl), a subspecies that has a large repertoire of GHs [23] and encodes a core-1 O-glycan degradation pathway [24–26]. Except EcN (Creative Biolabs, Shirley, NY), all strains were sourced from ATCC (Manassas, VA).

Media components included MRS broth (Sigma-Aldrich; used for recovery of *Bifidobacteria* and *Lactobacilli*) and YCFAC (Anaerobe Systems, Morgan Hill, CA), a complex medium composed of casitone, glucose, cellobiose, maltose, volatile fatty acids, vitamins, salts, hemin, and resazurin (final pH = 6.8± 0.1). Natural mucins used as culture medium supplements were porcine gastric mucin (PGM, Type II; Sigma-Aldrich) and chemically purified Muc2 isolated from porcine intestine (Muc2, gift of Ribbeck laboratory at MIT). Synthetic mucin mimetics were N-acetylglucosamine (GlcNAc)- or N-acetylgalatosamine (GalNAc)-substituted *Bombyx mori* silk proteins in which monosaccharides were conjugated to serine and threonine residues on a silk protein backbone. The synthesis of GlcNAc- and GalNAc-modified silk proteins followed the step-wise protocol described previously [27]. All culturing was conducted in a vinyl anaerobic chamber (Anaerobe Systems AS150) maintained under a 5% H_2_, 5% CO_2_, 90% N_2_ atmosphere (part no. ZNZ03NI90N2003194; Airgas, Radnor, PA). Reagents, consumables, and media were introduced into the chamber 24 h before use to ensure anaerobic equilibration.

### Anaerobic Culture

For all experiments, bacterial strains were outgrown from glycerol stocks onto pre-reduced agar plates and into 1 ml starter cultures in the appropriate recovery medium (100% MRS for *Bifidobacteria* and *Lactobacilli*, and 100% YCFAC for all other strains), and incubated for 24 h at 37°C. This initial growth stage was used to revive cells from cryostorage. Cultures were then pelleted (2,900×*g*, 10 min), washed in sterile 1× PBS, and transferred into 1 ml of 100% YCFAC for a second 24 h incubation to promote outgrowth and media adaptation. During these two stages, Am was further supplemented with 0.1% (w/v) autoclaved PGM to allow adequate growth. All washing and transfers were performed within the anaerobic chamber, while centrifugation was performed externally in anaerobically-sealed plates. Biological replicates were generated from independent agar plate colonies when feasible, or from separately inoculated tubes prepared from distinct glycerol stocks in the case of Am, which does not show robust growth on YCFAC agar plates. Cell-free media controls were included at each stage to monitor for contamination, and base YCFAC medium inoculated with each strain served as a substrate-independent growth control for cell viability.

Following the two-stage pre-growth procedure, cultures were inoculated at a starting OD_600_ of 0.05 ±0.01 in a final volume of 400 µL into 96-well deep-well plates containing 25% YCFAC in 1× PBS supplemented with 0.2% (w/v) of PGM, Muc2, SA, or SU. Cultures were incubated at 37°C for 24 h inside the anaerobic chamber. At the end of the incubation, contents were transferred to a clear 96-well plate and OD_600_ values were recorded. The plates were then centrifuged (2,900 *g*, × 10 min), and supernatants and cell pellets were separated and immediately frozen at –80°C. All conditions were tested in two independent experiments, each including biological and technical duplicates.Each experiment included a cell-free control (no inoculum) and a base YCFAC medium control (inoculated but without mucin supplementation).

### Sugar extraction and PMP-derivitization

Prior to LC-MS analysis, sugars were extracted from supernatant samples and derivatized using 3-Methyl-1-phenyl-2-pyrazoline-5-one (PMP). After thawing on ice, 25 µL of each culture supernatant sample was mixed with 5 µL of internal standard (D-Glucose-^13^*C*_6_, 25 µg/mL), followed by the addition of 60 µL of ice-chilled methanol.

After vortexing and centrifuging at 4000 rpm for 10 minutes, 60 µL of each supernatant mixture was transferred to a well of a new plate. Then, 20 µL of freshly prepared 0.5M PMP solution (87 mg/mL in methanol) and 16.6 µL of ammonium hydroxide solution (20% in water) were added, followed by vortexing. The mixture was incubated at 60°C for 30 min to facilitate the derivatization reaction. The reaction was quenched with 16.6 µL of formic acid, after which the mixture was vortexed and briefly spun down. After sequential extractions with chloroform, 40 µL of the final aqueous phase was collected and diluted with 120 µL of HPLC-grade water prior to LC-MS injection.

### Targeted LC-MS analysis of PMP-derivatized sugars

Chromatographic separation of PMP-derivatized sugars was performed on a Gemini 5 µm C18 110 Å column (250 × 2 mm; Phenomenex, Torrance, CA) using a gradient elution method. The LC-MS system was an 1260 Infinity III LC System (Agilent, Santa Clara, CA) coupled to a TripleTOF 5600 quadrupole-time-of-flight (QToF) mass spectrometer (AB Sciex, Framingham, MA). The mobile phases consisted of solvent A (acetonitrile:water, 5:95 v/v) with 20 mM ammonium formate (pH adjusted to 9.45) and solvent B (100% acetonitrile). The flow rate was set to 450 µL/min, with an injection volume of 10 µL. The column oven temperature was maintained at 55 °C. The elution gradient started at 90% A, decreased to 80% by 2 min, to 75% A at 10 min, and to 5% A at 12 min. The solvent composition was held at 5% A until 20 min, then returned to 90% A by 20.5 min and maintained until 28 min. The mass spectrometer was operated in negative electrospray ionization mode. Data acquisition was performed using multiple product ion experiments (Table 1).

**Table 1.** LC-MS parameters for targeted analysis of mucin O-glycan sugars.

| Sugar | Precursor<br>(m/z) | CE<br>(eV) | DP<br>(V) | 1<sup>st</sup> Product<br>(m/z) | 2<sup>nd</sup> Product<br>(m/z) | RT<br>(min) | R<sup>2</sup> |
| --- | --- | --- | --- | --- | --- | --- | --- |
| D-Mannose | 509.20 | -17 | -90 | 215.08 | 335.12 | 5.24 | 0.993 |
| D-Glucose | 509.20 | -17 | -90 | 215.08 | 335.12 | 7.53 | 0.994 |
| D-Galactose | 509.20 | -17 | -90 | 215.08 | 335.12 | 7.97 | 0.993 |
| L-Rhamnose | 493.21 | -20 | -120 | 215.08 | 319.13 | 5.91 | 0.995 |
| D-Fucose | 493.21 | -20 | -120 | 215.08 | 319.13 | 8.63 | 0.978 |
| MalNAc | 550.23 | -30 | -120 | 256.11 | 376.15 | 8.40 | 0.996 |
| GalNAc | 550.23 | -30 | -120 | 256.11 | 376.15 | 10.01 | 0.990 |
| GlcNAc | 550.23 | -30 | -120 | 256.11 | 376.15 | 8.05 | 0.996 |
| Neu5Ac | 446.16 | -30 | -120 | 173.07 | 297.10 | 2.67 | 0.990 |
| D-Glucose-<sup>13</sup>C<sub>6</sub> | 515.22 | -17 | -90 | 217.09 | 341.14 | 7.53 | IS |

Calibration curves were constructed by linear regression of areas-under-the-curve for pure chemical standards, with standard concentrations ranging from 7–500 $\mu$ M. CE: collision energy; DP: declustering potential; RT: retention time; IS: internal standard.

## Results

### DEFT provides superior enzyme classification performance

We benchmarked DEFT against four state-of-the-art sequenced-based methods for EC number prediction: ECPred [7], DeepEC [5], ProteInfer [4], and CLEAN [6]. We also compared DEFT’s performance against a naive structure-based approach that transfers the entire EC number from an already annotated enzyme that is structurally most similar to the enzyme of interest. **ECPred** takes an ensemble approach to EC number prediction by combining multiple feature types. It integrates subsequence-level features, homology-based information, and biochemical properties into a weighted scoring scheme for final predictions. While effective for well-characterized enzyme families, ECPred’s reliance on pre-computed features limits its ability to generalize to novel enzyme architectures. **DeepEC** pioneered the application of deep learning to enzyme classification. The method represents protein sequences as one-hot encoded matrices of size ℝ^*m×*21^(where *m* is sequence length and 21 accounts for the 20 amino acids plus unknown residues) and applies convolutional neural networks. DeepEC employs a multi-task architecture with three separate convolutional layers: one for binary enzyme/non-enzyme classification, and two others for predicting EC levels 1-3 and level 4, respectively, in a multilabel fashion. **ProtInfer** builds upon DeepEC’s convolutional approach but incorporates dilated convolutions to capture long-range sequence dependencies. By gradually increasing the receptive field size, ProtInfer can better model the relationship between distant residues that may be critical for enzymatic function. **CLEAN** represents the current state-of-the-art in enzyme classification. It employs a contrastive learning framework that embeds protein sequences using ESM-1b followed by LayerNorm. The method learns representations where enzymes with similar EC annotations are brought closer together in embedding space while dissimilar enzymes are pushed apart. This approach has shown superior performance on standard benchmarks and provides well-curated evaluation datasets with controlled sequence similarity.

We used **Foldseek** [15] to detect candidate enzymes for structure-based alignment and EC number annotation. Briefly, the enzyme of interest’s 3D structure is first discretized with Foldseek’s tokenizer yielding a 3Di string, which describes the geometric conformation of each residue *i* in the protein backbone with its spatially closest residue *j*. The 3Di string is then aligned against our reference set, and the best match (as reported by E-values) has its EC number transferred to the unknown enzyme.In our benchmarking experiments we chose our reference set to be the enzymes in the training split.

The benchmarking experiments followed the framework of the CLEAN study by Yu et al. [6]. The same two datasets, New-392 and Price-149, were used, with the same training/testing splits. The datasets comprise, respectively, 392 new additions to UniProt introduced after the curation of the SwissProt datasets and 149 proteins released by ProteInfer [4] as historically difficult to annotate. To afford comparisons with the previous study by Yu et al., we also used the original SwissProt splits from CLEAN for hyperparameter selection, taking care to ensure that there is no overlap between the training and evaluation sets. On both New-392 and Price-149 datasets, DEFT showed superior precision and recall performance compared with all other methods as measured by the F1 score. Specifically on Price-149, DEFT achieved an F1 score of 0.72 compared with 0.48 for CLEAN, the next best method (Fig. 2a). On New-392, DEFT achieved an F1 score of 0.84 compared with 0.50 for CLEAN (Fig.2b). Overall, the performance improvements by DEFT ranged from 1.5-(compared with CLEAN on Price-149) to 36-fold (compared with ECPred on Price-149).

**Fig 2.**
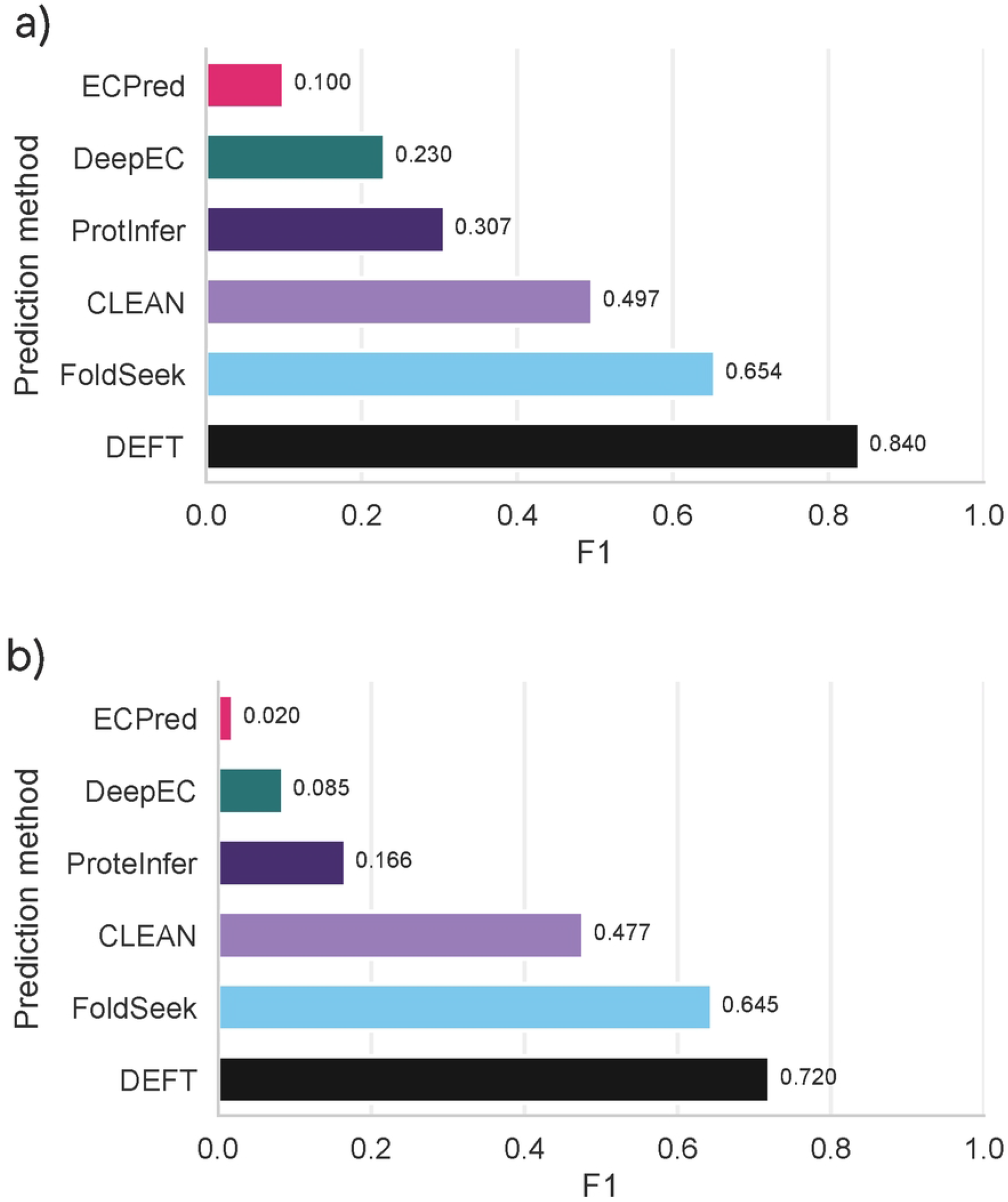
DEFT classification performance. Classification accuracy (F1 score) of DEFT compared with existing EC prediction methods on the New-392 (a) and Price-149 (b) datasets. The F1 score is the harmonic mean of precision and recall.

We further analyzed the cross-validation results to characterize the dependence of annotation accuracy on the frequency of an EC number in the training dataset. Table 2 displays the precision and recall of DEFT stratified by number of times a particular EC number occurs in the training dataset. The precision and recall metrics shown in Table 2 compare DEFT’s cross-validation performance on the same SwissProt dataset with at most 50% sequence similarity between any two proteins. As expected, precision and recall of both DEFT and CLEAN declined as the frequency of a particular EC number in the training dataset decreased. However, the performance of CLEAN declined more sharply. Even for rare EC numbers (less than 5 occurrences), DEFT’s recall was 0.87 ± 0.03, whereas CLEAN’s recall declined to 0.69 *±* 0.03.

**Table 2.** Precision and recall of DEFT’s enzyme classifications on CLEAN’s 50% homology training split grouped by number of times the EC number is present in the training dataset.

| EC number occurrence | Precision |  | Recall |  |
| --- | --- | --- | --- | --- |
|  | CLEAN | DEFT | CLEAN | DEFT |
| 0–5 | $0.72 \pm 0.03$ | $0.75 \pm 0.01$ | $0.69 \pm 0.03$ | $0.87 \pm 0.03$ |
| 6–10 | $0.93 \pm 0.02$ | $0.94 \pm 0.03$ | $0.87 \pm 0.01$ | $0.95 \pm 0.02$ |
| 11–50 | $0.96 \pm 0.01$ | $0.98 \pm 0.01$ | $0.90 \pm 0.01$ | $0.99 \pm 0.01$ |
| 51–100 | $0.98 \pm 0.01$ | $1.00 \pm 0.01$ | $0.94 \pm 0.01$ | $0.99 \pm 0.01$ |
| 100–∞ | $0.97 \pm 0.00$ | $0.99 \pm 0.01$ | $0.92 \pm 0.01$ | $0.99 \pm 0.01$ |

### DEFT facilitates genome-wide profiling of enzyme repertoires

A key problem in biology is searching for homologous proteins. Typical techniques to search for homologous proteins involve BLAST like queries on the primary amino acid sequence against a reference database. Foldseek intuitively extends a BLAST like search to also utilize structural information, while maintaining the computational benefits of utilizing character searching algorithms. Often, the purpose of such a search is to find similarly functioning proteins in other species. The challenge with such a search in the context of enzymes is that through evolutionary pressure catalytic sites tend to be more conserved than the rest of the protein. In such a case, the regions of two proteins that need to be aligned tend to be small portions of the proteins. As a result, the signal of finding similar enzymes on the basis of global alignment tends to be drowned out by noise.

We tested DEFT’s capability to perform genome-wide enzyme classification by characterizing the GH profiles of several gut bacteria. The bacteria were selected to represent anaerobes that inhabit the intestinal mucus in mammals, and included both mucin grazers (Am and Bt) and non-grazers (Lp, Lr, Bd, and EcN). The analysis also included a Bl strain that has a large repertoire of GHs [23] but has been shown to poorly degrade mucin O-glycans [28, 29]. The inputs to the DEFT predictions were whole genomes of the bacteria and an expertly curated list of EC numbers representative of GH activities required for mucin O-glycan metabolism (Fig. 3).

**Fig 3.**
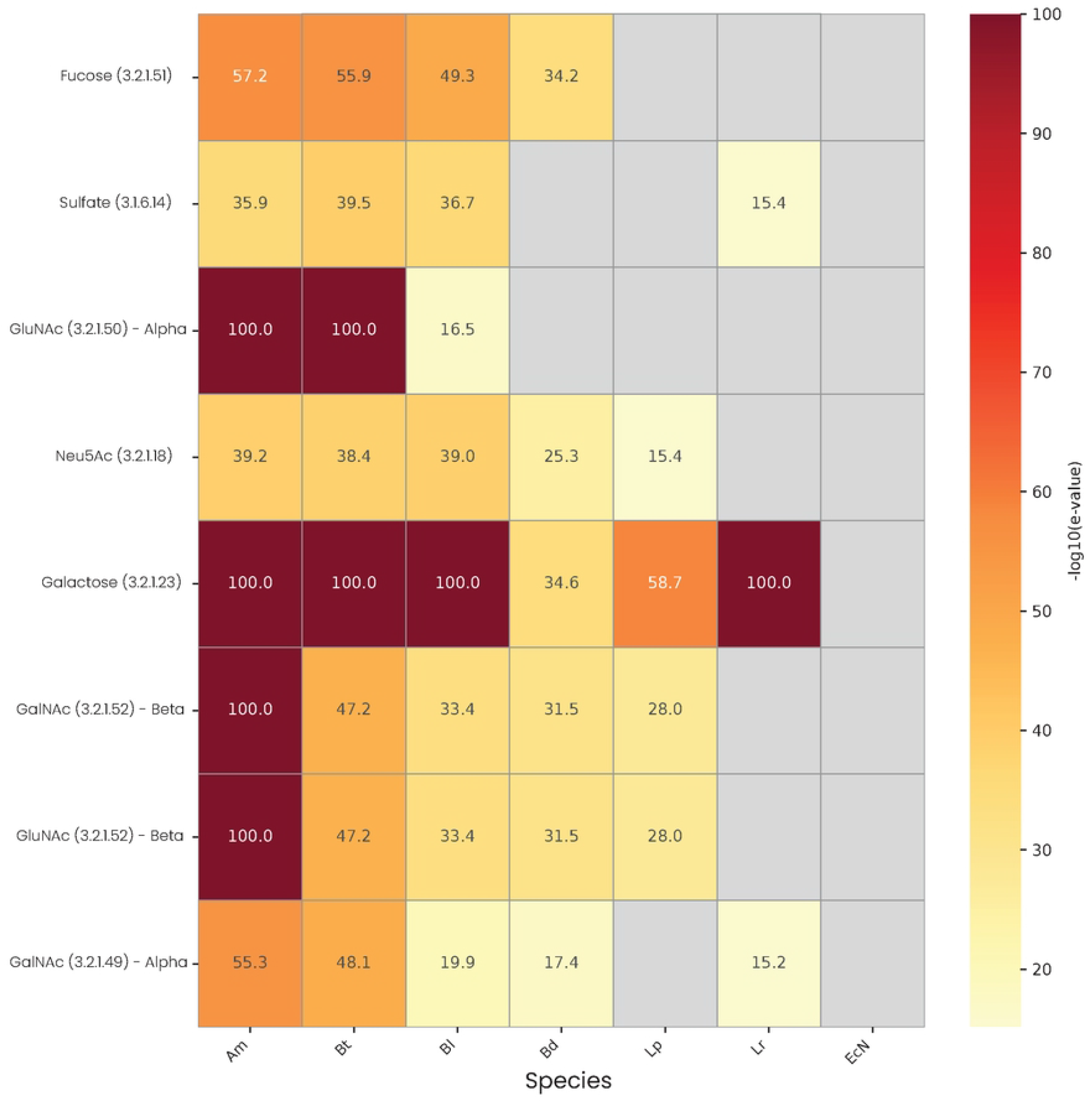
Matches for mucin O-glycan metabolizing enzymes in representative mucin-grazing and non-grazing gut anaerobes. The probability DEFT found a matching enzyme for an EC number of interest by chance is expressed by an E-value. See Methods for E-value definition. A scaling of− log_10_ is applied to emphasize the strength of the match. A gray squares indicates that DEFT did not find any matching enzymes in the organism for the EC number (see text for abbreviations of organism names). E-values less than 1 × 10^−60^ were truncated to -log values of 100

Genome-wide scans using DEFT found high-probability (low E-value) matches for all of the key GHs in the mucin grazers, including alpha-fucosidase (EC number 3.2.1.51), alpha-N-acetylgalactosaminidase (3.2.1.49), beta-N-acetylhexosaminidase (3.2.1.52), alpha-acetylglucosaminidase (3.2.1.50) and neuraminidase (3.2.1.18). In contrast, these enzymatic functions were either not detected in the non-grazers or matched with weaker (orders of magnitude higher) E-values. Interestingly, the predicted GH profile varied among the mucin grazers, with Am having stronger (orders of magnitude lower) E-values than Bt for hydrolysis of terminal N-acetylhexosamines from glycan chains (EC number 3.2.1.52).

### Experimental growth rates and sugar measurements validate DEFT predicted enzyme profiles of mucin-grazing and non-grazing bacteria

We experimentally evaluated the predicted ability of selected gut bacteria to utilize mucin O-glycans as metabolic substrates by measuring their growth on various mucin supplemented culture media. The substrates used for the experiments were porcine gastric mucin (PGM), porcine mucin 2 (Muc2), and two mucin mimetics that have simpler, defined O-glycan profiles. The mimetics were *Bombyx mori* silk proteins modified with GalNAc (SA) or GlcNAc (SU) at the hydroxyl groups of serine and threonine (S/T) residue derivatives. Without mucin supplementation, bacterial growth did not stratify [27] by the ability to metabolize mucin O-glycans (Fig. 4). Growth was slowest for Am, reaching an *OD*_600_ value less than 0.1. The fastest growing group comprised Bt, Lp, Lr, and Bd, which reached *OD*_600_ values between 0.35-0.45. Mucin supplementation had the largest growth promoting effect on Am. This is consistent with the predicted GH profiles (Fig. 3) showing Am with the strongest E-values for enzyme functions (EC numbers) needed to hydrolyze common glycosidic bonds in mucin O-glycans [30]. With PGM or Muc2 supplementation, Am growth was 4-fold higher compared with the base YCFAC medium. Supplementation with SA or SU had a weaker, but still significant (∼36 to ∼41%) growth-promoting effect for Am. Mucin supplementation also enhanced the growth of Bt, which was predicted to encode a GH profile comparable to Am, albeit with a weaker E-value for EC number 3.2.1.52. The increases in Bt growth compared with the base medium ranged from ∼15 (SU) to∼24% (Muc2). Mucin supplementation had no significant effect on growth of Bl, which had weaker E-values than Am or Bt for EC numbers (3.2.1.49, 3.2.1.52, and 3.2.1.50) corresponding to release of N-acetylhexosamines (Fig. 3). As expected, the other non-grazers (Bd, Lp, Lr, and EcN) did not respond significantly to mucin supplementation.

**Fig 4.**
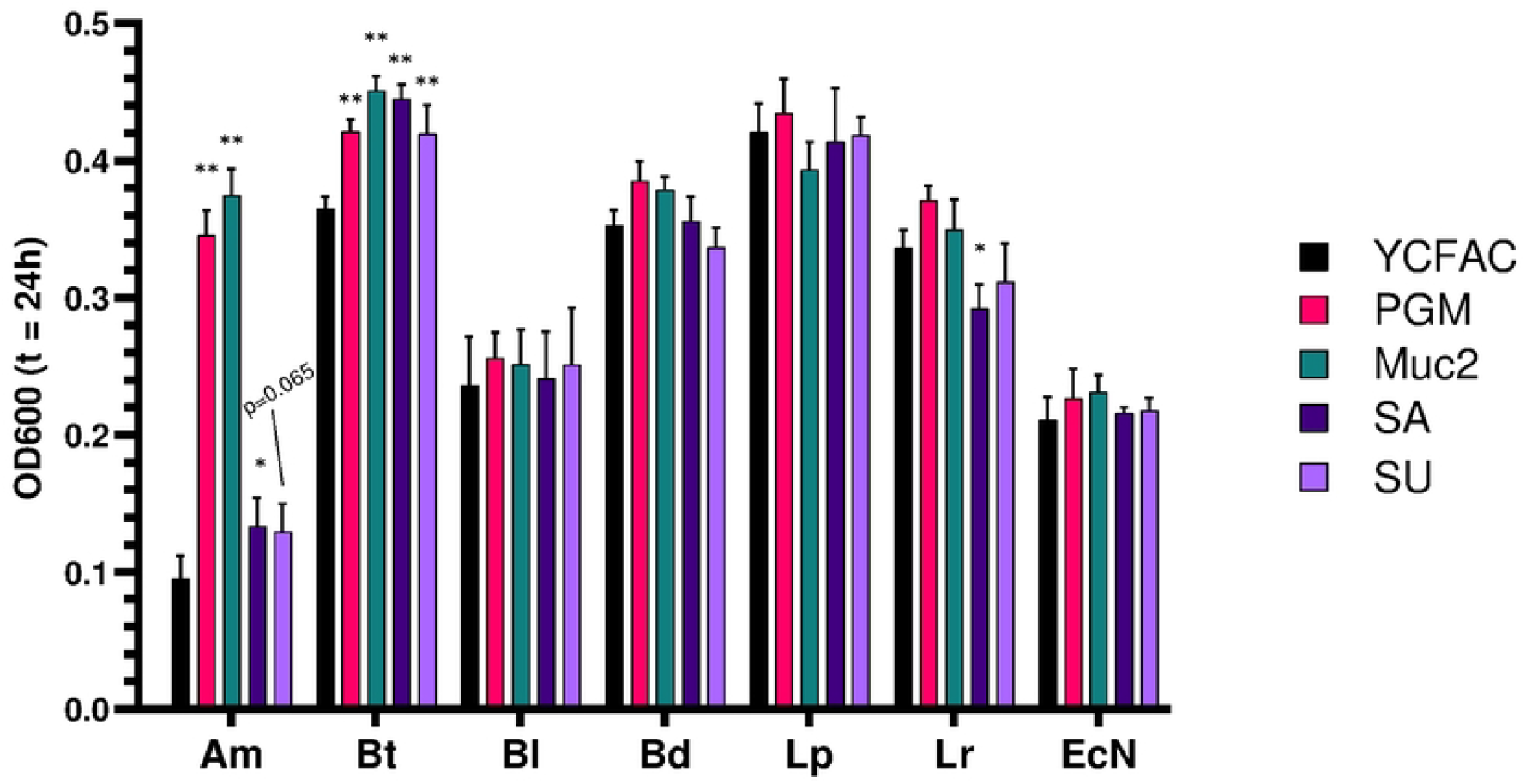
Anaerobic growth of mucin grazers and non-grazers in mucin supplemented media. All cultures were inoculated at an approximate *OD*_600_ value of 0.05. Final *OD*_600_ values were recorded after 24 h of culture in base (YCFAC) or mucin-supplemented medium (PGM, Muc2, SA, or SU). Data shown are mean± SD of n = 4 biological replicates. Statistical significance was determined using two-way ANOVA followed by multiple comparisons with Dunnett’s adjustment relative to the YCFAC, non-mucin control. ^*\**^ p*<* 0.05; ^*\*\**^ p*<* 0.01.

To further investigate the differential utilization of mucins by the bacteria, we measured the medium concentrations of sugars comprising the bulk of O-glycan core structures and termini. Targeted LC-MS assays showed significant increases in GalNAc, GlcNAc, N-acetylneuraminic acid (Neu5Ac), fucose, and galactose in Muc2-supplemented Am cultures (Fig. 5a-e). Except for fucose, the other four sugars were also significantly increased in Am cultures. However, the increases in GlcNAc and Neu5Ac were lower compared with Am cultures with Muc2 supplementation.

**Fig 5.**
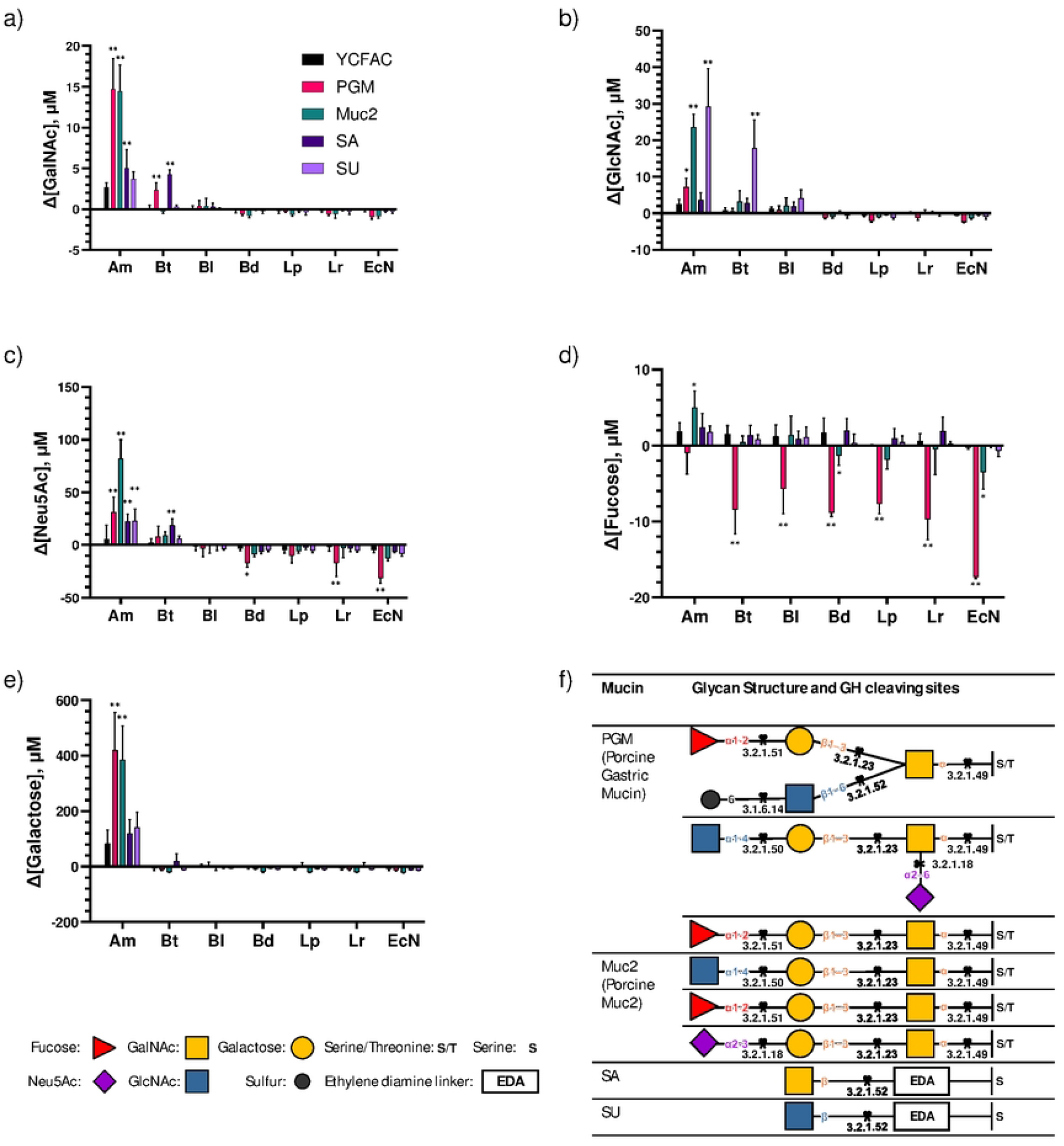
Changes in O-glycan core and terminal sugar concentrations. (a–e) Changes in O-glycan core and terminal sugar concentrations following incubation of mucin-grazing and non-grazing gut bacteria in base YCFAC medium or YCFAC supplemented with mucin substrates (PGM, Muc2, SA, or SU). See text for sugar name abbreviations. Bars represent medium (cell-free) control subtracted concentrations.Positive values indicate net accumulation of monosaccharide; negative values indicate net consumption. Statistical significance was determined using two-way ANOVA followed by multiple comparisons with Dunnett’s adjustment relative to the YCFAC, non-mucin control. Data shown are mean ± SD of n = 2 biological replicates from N = 2 independent experiments. ^*\**^ p*<* 0.05; ^*\*\**^ p*<* 0.01. Table (f) shows putative top three O-glycan structures present in each mucin substrate (PGM [31], Muc2 [32], SA, or SU) and the potential EC numbers of GHs required to cleave the bonds.

Incubation of Am in SA- or SU-supplemented medium selectively increased the concentrations of GalNAc and GlcNAc, respectively (Fig. 5a&b), consistent with predicted hydrolysis of the sugars from the mucin mimetics (Fig. 5f). Compared with Am, Bt cultures showed only limited increases in sugar concentrations when incubated with mucins. Incubation of Bt with PGM significantly increased GalNAc compared with the cell-free control, while incubation with Muc2 did not increase any measured sugars. As was the case for Am, incubation of Bt in SA- or SU-supplemented medium selectively increased the concentrations of GalNAc and GlcNAc, respectively (Fig. 5a&b). Besides Am and Bt, no other bacteria increased any measured sugars.

Interestingly, Neu5Ac levels were significantly reduced relative to the cell-free control when Bd, Lr, or EcN were incubated in PGM-supplemented medium (Fig. 5c). Except for Am, all other bacteria exhibited significant net consumption of fucose when incubated in PGM-supplemented medium (Fig. 5d). As these trends were not observed when the bacteria were incubated in Muc2-supplemented medium, we investigated if PGM- and Muc2-supplemented fresh media had different sugar profiles. Targeted LC-MS analysis revealed that the PGM-supplemented medium, without exposure to cells, had significantly elevated levels of Neu5Ac and fucose compared with the base YCFAC medium (S1 Fig a). This suggested that fresh PGM-supplemented medium may contain free sugars from non-enzymatic degradation of glycan termini during sterilization or sample preparation. To test this possibility, we measured sugar concentrations in different dilutions of PGM (0.05-0.8% w/v). This analysis found that the sugar concentrations inversely correlated with PGM dilution (S1 Fig b). Fucose and Neu5Ac were detected at the highest levels (∼60 *µ*M at 0.8% w/v PGM). These results confirmed that PGM supplementation introduced significant levels of free sugars to the base medium, independent of bacterial activity. However, at the level of PGM supplementation used in this study (0.2% w/v), only fucose and Neu5Ac were present as free sugars at sufficient concentrations to support significant consumption by the cells (Fig. 5).

The profiles of non-glycan sugars qualitatively differed from those of glycan sugars (Fig. 6). Significant increases in ManNAc were measured in SU- and SA-supplemented Am cultures (1.6-to 2.1-fold higher, respectively, than Am culture in base medium without mucin) (Fig. 6a). However, ManNAc was not used to modify either silk protein. Mannose levels of mucin-supplemented Am and Bt cultures did not change significantly compared with the corresponding base medium controls (Fig. 6b). Both Bl and Lr cultures produced mannose when incubated in SU- or SA-supplemented medium, whereas Bd cultures consumed mannose when incubated in PGM-, Muc2-, or SU-supplemented medium. Glucose, present as a component of the base medium (S1 Fig a), was substantially depleted by Bt, Bl, Bd, Lp, Lr, and EcN under all medium conditions, indicating a shared reliance on free glucose as a carbon source. By comparison, Am only depleted glucose in PGM- or Muc2-supplemented medium. Together with the lower OD_600_ value of Am in YCFAC and SA- or SU-supplemented medium, these results suggest that basal glucose consumption correlates with cell growth. Overall, the measured bacterial profiles of non-glycan sugars, unlike glycan sugars, did not cluster according to the predicted GH repertoire.

**Fig 6.**
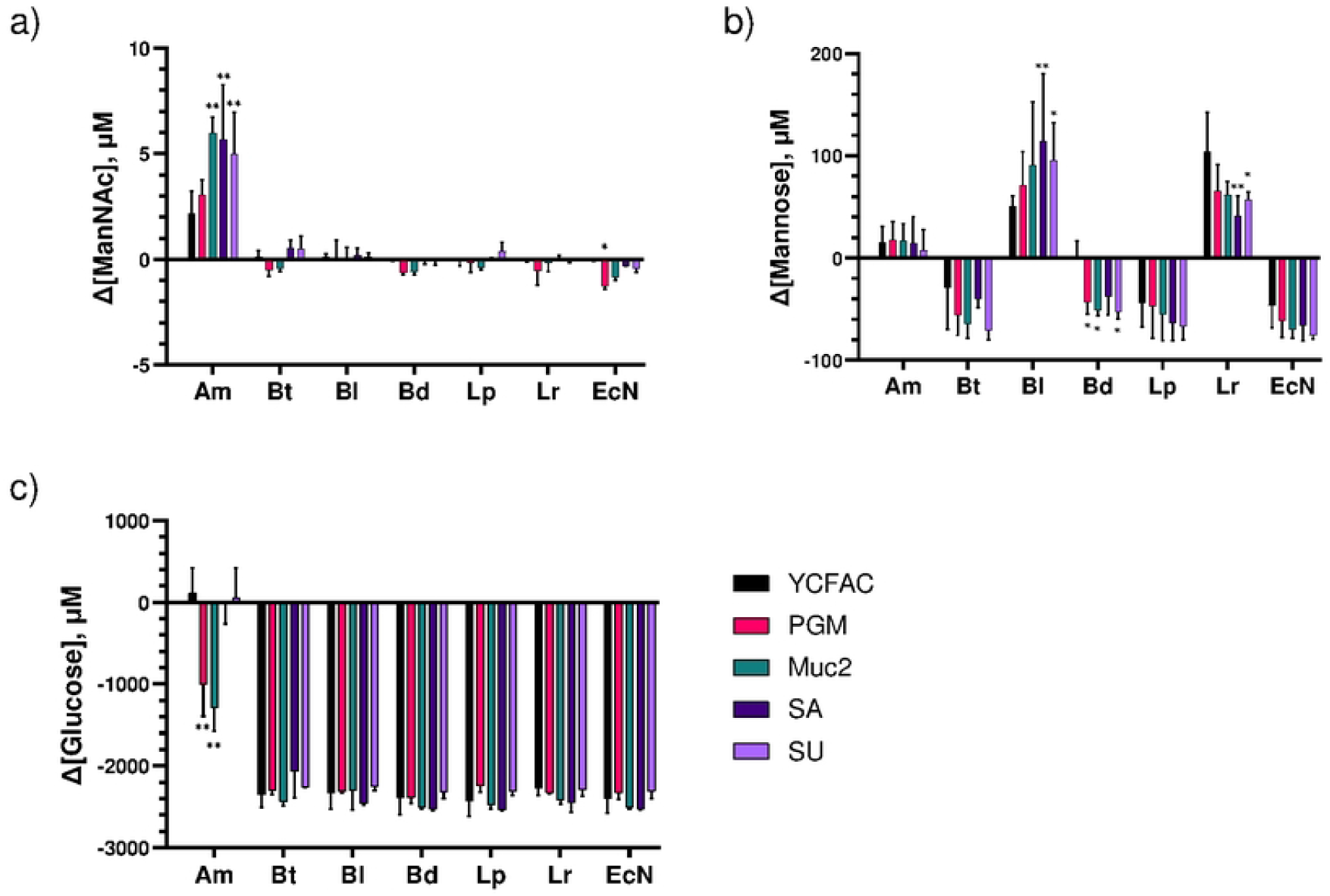
Changes in non-mucin sugar concentrations. (a–c) Changes in non-glycan sugar concentrations following incubation of mucin-grazing and non-grazing gut bacteria in base YCFAC medium or YCFAC supplemented with mucin substrates (PGM, Muc2, SA, or SU). See text for sugar name abbreviations. Bars represent medium (cell-free) control subtracted concentrations. Positive values indicate net accumulation of monosaccharide; negative values indicate net consumption. Statistical significance was determined using two-way ANOVA followed by multiple comparisons with Dunnett’s adjustment relative to the YCFAC, non-mucin control. Data shown are mean ± SD of n = 2 biological replicates from N = 2 independent experiments. ^*\**^ p*<* 0.05; ^*\*\**^ p*<* 0.01..

## Discussion

The present study introduces a novel enzyme classification method, DEFT, which takes a hybrid of two different strategies on coarse and fine prediction of EC number to substantially outperform existing methods. Applying the same benchmarking approach as the study by Yu et al. [6], we demonstrate that DEFT achieves substantial improvements in precision and recall, achieving F1 scores 1.7- and 1.5-fold higher than the next best performing method (CLEAN) on New-392 and Price-149 datasets, respectively. The hybrid strategy takes advantage of the fast structural search capabilities of Foldseek [15] to perform the fine classification of the last two EC number digits, while avoiding the pitfall of using structural alignment to assign the most general EC number levels (first two digits), instead using the SaProt PLM representation to learn the correct portions of the structure that are important for enzymatic activity.

SaProt-based assignment of the first two EC number levels, followed by Foldseek structural search-based assignment of the last two EC number levels, is very fast.Annotating 5,000 proteins takes under 5 minutes on a single NVIDIA H200 machine. This enables DEFT to be used for genome-wide profiling of an organism’s entire enzyme repertoire. As an illustrative use case, we used DEFT to predict the ability of representative gut bacteria to hydrolyze mucin O-glycans. The predictions were validated experimentally by analyzing bacterial growth and glycan sugar level changes in mucin-supplemented media. The experimental results independently verified the predicted mucin-grazing enzyme profiles of Am and Bt. DEFT also correctly predicted the non-grazing enzyme profiles of the remaining species. These results are in good agreement with previous studies reporting that Am and Bt can degrade mucin O-glycans and utilize the glycan sugars as substrates for growth [33, 34].

Although Bl is a potential mucin grazer because it has an endo-*α*-N-acetylgalactosaminidase (engBF) [25] and an intracellular degradation pathway for core-1 structure (Gal*β*1-3GalNAc) [24, 26], the engBF encoded endo-*α*-N-acetylgalactosaminidase is substrate-restricted to act only on the core-1 structure [25], which is rare in natural mucins. A previous study reported that among four *Bifidobacterium longum* subsp. *longum* strains encoding engBF genes, only one strain (NCIMB8809) exhibited appreciable mucin O-glycan degradation activity [29]. Combined with the lack of a comprehensive O-glycan degradation enzyme repertoire in Bl—as revealed by DEFT’s genome—wide analysis-this likely explains why *Bifidobacterium longum* subsp. *longum* ATCC 15707 is unable to hydrolyze PGM or Muc2 glycans into monosaccharides. Using gut bacterial mucin O-glycan degradation as a case study, we demonstrate DEFT’s strong ability to infer complex organismic functions by predicting multiple catalytic activities (EC numbers) in a computationally efficient manner.

We note that a correct EC number is sometimes not a sufficiently fine specification of enzyme activity. Taking glycan chain degradation as an example, current EC number assignments do not clearly distinguish among functionally distinct activities. A representative case is EC number 3.2.1.52 [35, 36], assigned to *β*-N-acetylhexosaminidase. This enzyme has been reported to exhibit both endo- and exo-acting GH activities [37], as well as activity toward both GalNAc and GlcNAc. However, endo-acting GHs can, in some contexts, degrade mucins more effectively because they initiate cleavage within the oligosaccharide chain rather than acting only at the termini. Whether DEFT, provided with appropriate training examples, can capture such subtle yet critical activity distinctions warrants further study. Prospectively, users may provide a curated enzyme training set with pseudo–last (e.g., fifth) EC number digits encoding user-defined, context-dependent activity subclasses. Without retraining the entire enzyme classification model, DEFT should in principle be able to learn structural similarities and make predictions using the user-provided protein sequences along with structural information and custom subclass definitions. In this way, DEFT has the potential to bridge the gap between rigid EC number classification and the functional complexity inherent in biological systems, a challenge commonly encountered in studying biological degradation of both natural and synthetic polymers.

## Data and software availability

The code for DEFT is available at https://github.com/merterden98/DEFT Model training weights and data are archived at https://zenodo.org/records/17858733

## Supporting information

**S1 Fig. Free sugar concentrations in fresh media and PGM solutions**. Free sugar concentrations in (a) mucin-supplemented fresh (cell-free) media and (b) different PGM dilutions in 1x PBS (0.05-0.8% w/v). The concentrations were determined using targeted LC-MS analysis as described in Methods.

## Acknowledgments

We thank the Tufts BCB group for helpful discussions and the Ribbeck lab for the purified mucin.

